# Histone H3K36me2-specific methyltransferase ASH1L is required for the MLL-AF9-induced leukemogenesis

**DOI:** 10.1101/2021.08.05.455280

**Authors:** Mohammad B. Aljazi, Yuen Gao, Yan Wu, George I Mias, Jin He

**Author notes:** Correspondence: Jin He.

## Abstract

ASH1L and MLL1 are two histone methyltransferases that facilitate transcriptional activation during normal development. However, the roles of ASH1L and its enzymatic activity in the development of MLL-rearranged leukemias are not fully elucidated in the *Ash1L* gene knockout animal models. In this study, we used an *Ash1L* conditional knockout mouse model to show that loss of ASH1L in hematopoietic progenitor cells impaired the initiation of MLL-AF9-induced leukemic transformation *in vitro*. Furthermore, genetic deletion of ASH1L in the MLL-AF9-transformed cells impaired the maintenance of leukemic cells *in vitro* and largely blocked the leukemia progression *in vivo*. Importantly, the loss of ASH1L function in the *Ash1L*-deleted cells could be rescued by wild-type but not the catalytic-dead mutant ASH1L, suggesting the enzymatic activity of ASH1L was required for its function in promoting MLL-AF9-induced leukemic transformation. At the molecular level, ASH1L enhanced the MLL-AF9 target gene expression by directly binding to the gene promoters and modifying the local histone H3K36me2 levels. Thus, our study revealed the critical functions of ASH1L in promoting the MLL-AF9-induced leukemogenesis, which provides a molecular basis for targeting ASH1L and its enzymatic activity to treat MLL-arranged leukemias.

## Introduction

The MLL rearrangement (MLLr) caused by 11q23 chromosomal translocations creates a variety MLL fusion proteins that drive the acute lymphoblastic and myeloid leukemia development, which accounts for approximate 5-10% acute leukemias in human patients[1–5]. Despite recent progression in the development of chemotherapies against leukemias, the overall prognosis for the MLLr leukemias remains poor[6, 7].

MLL1 protein is a histone lysine methyltransferase (KMTase) that contains a SET (*<u>S</u>*u(var)3-9, *<u>E</u>*nhancer-of-zeste and *<u>T</u>*rithorax) domain to catalyze trimethylation of histone H3 lysine 4 (H3K4me3)[8]. Functionally, MLL1 belongs to the Trithorax-group (TrxG) proteins that antagonize the Polycomb-group (PcG)-mediated gene silencing and facilitate transcriptional activation[9]. In 11q23 chromosomal translocations, the *N*-terminal portion of MLL1 is fused with a variety of fusion partners to generate different oncogenic MLL fusion proteins that function as disease drivers leading to leukemia development[10–12]. Previous studies have revealed that the *N*-terminal portion of MLL fusion proteins interacts with MENIN and LEDGF (*<u>L</u>*ens *<u>E</u>*pithelium-*<u>D</u>*erived *<u>G</u>*rowth *<u>F</u>*actor), which is critical for the recruitment of MLL fusion proteins to chromatin, whereas the *C*-terminal fusion partners interact with various trans-activators to induce transcriptional activation[13–17]. However, since the MLL fusion proteins lack the intrinsic histone H3K4 methyltransferase activity due to loss of SET domain located in the *C*-terminal portion of MLL1[10], it is unclear whether other histone modifications are required for the MLL fusion proteins-induced gene expression and leukemogenesis.

Recently, another member of TrxG proteins, ASH1L (*<u>A</u>*bsent, *<u>S</u>*mall, or *<u>H</u>*omeotic-*<u>L</u>*ike *<u>1</u>*), was found to play important roles in normal hematopoiesis and leukemogenesis[8, 18, 19]. Biochemically, ASH1L is a histone KMTase that mediates dimethylation of histone H3 lysine 36 (H3K36me2)[20]. Similar to MLL1, ASH1L facilitates gene expression through antagonizing PcG-mediated gene silencing[8]. Previous studies have shown that ASH1L and MLL1 co-occupies the same transcriptional regulatory regions, and loss of either ASH1L or MLL1 reduces the expression of common genes[21–23], suggesting ASH1L and MLL1 function synergistically to activate gene expression during normal development. However, the significance of ASH1L and its-mediated histone H3K36me2 in the MLLr-associated leukemogenesis has not been addressed in the *Ash1L* gene knockout animal models.

In this study, we used an *Ash1L* conditional knockout mouse model to show that loss of ASH1L in hematopoietic progenitor cells (HPCs) impaired the initiation of MLL-AF9-induced leukemic transformation *in vitro*. Furthermore, genetic deletion of ASH1L in the MLL-AF9-transformed cells impaired the maintenance of leukemic cells *in vitro* and largely blocked the leukemia progression *in vivo*. Importantly, the loss of ASH1L function in the *Ash1L*-deleted cells could be rescued by ectopic expression of wild-type but not the catalytic-dead mutant ASH1L, suggesting the enzymatic activity of ASH1L was required for its function in promoting MLL-AF9-induced leukemic transformation. At the molecular level, ASH1L activated the MLL-AF9 target gene expression by directly binding to the gene promoters and modifying the local histone H3K36me2 levels. Thus, our study revealed the critical functions of ASH1L in MLL-AF9-induced leukemogenesis and raised the possibility that ASH1L might serve as a potential therapeutic target for the treatment of MLLr leukemias.

## Materials and Methods

### Mice

The *Ash1L* conditional knockout mice were generated as previously reported[24]. To generate inducible *Ash1L* deletion, mice were crossed with Rosa26-CreER^T2^ mice that were obtained from The Jackson Laboratory. All mice for this study were backcrossed to C57BL/6 mice for at least five generations to reach pure genetic background prior to conducting experiments. All mouse experiments were performed with the approval of the Michigan State University Institutional Animal Care & Use Committee.

### Hematopoietic progenitor isolation, and culture

Hematopoietic progenitor cells were isolated from femurs of 4-to 6-week C57BL/6 mice. The red blood cells in the bone marrows were lysed by ammonium chloride solution (Stem Cell Technologies 07800) and filtered with a 70-μm nylon filter. The c-kit^+^ HPCs were isolated using c-Kit antibody-conjugated IMag (BD Biosciences) beads. Cultures of transformed HPCs cells were maintained in RMPI media supplemented with 10% FBS, 1% MEM non-essential amino acids, 1% Glutamax, 10 ng/mL, and 2-mercaptoethanol. To induce Cre-mediated recombination *in vitro*, 4-hydroxy-tamoxifen (Sigma-Aldrich) was resuspended in DMSO and supplemented into the culture medium with concentration of 250 nM.

### Retroviral and lentiviral vector production and transduction

The pMIG-FLAG-MLL-AF9 retroviral vectors as obtained from Addgene (Plasmid #71443). Retroviral vectors were generated by co-transfection of retroviral vectors with pGag-pol, pVSVG 293T cells using CalPhos mammalian transfection kit (TaKaRa). After 48hrs post transfection, viral supernatant was harvested, filtered through a 0.45 μm membrane, and concentrated by ultracentrifugation. The lentiviral system was obtained from the National Institutes of Health AIDS Research and Reference Reagent Program. To generate GFP expression vectors, the GFP cDNA was PCR amplified, fused with P2A and puromycin resistant cassette and cloned into the SpeI/EcoRI sites under the EF1α promoter. To generate lentiviral viruses, the transducing vectors pTY, pHP and pHEF1α–VSVG were co-transfected into HEK293T cells. The supernatant was collected at 24, 36 and 48 hours after transfection, filtered through a 0.45 μm membrane and concentrated by ultracentrifugation. Retroviral and lentiviral transduction of HPCs was performed by spin inoculation for 1 hour at 800g, in RMPI media supplemented with 10% FBS, 1x MEM non-essential amino acids (Life Technologies), 1x Glutamax (Life Technologies), 1x sodium pyruvate (Life Technologies), 50 ng/mL mSCF (PeproTech), 10 ng/mL mIL-6 (PeproTech), and 10 ng/mL mIL-3 (PeproTech).

### Serial methylcellulose replating assay and leukemia transplantation

The colony formation assays were conducted by plating 500 cells into methylcellulose media consisting of Iscove MDM (Life Technologies) supplemented with FBS, BSA, insulin-transferrin (Life Technologies), 2-mercaptoethanol, mSCF, mIL-3, mIL-6, and 10 ng/mL GM-CSF (PeproTech). After 7-10 days, the colony numbers were counted under a microscope. The colonies were picked up, and cells were pooled and replated onto secondary methylcellulose plates. Three rounds of replating were performed for each experiment. For leukemia transplantation, recipient C57BL/6 mice were subjected to total body irradiation at a dose of 11 Gy with the use of a X-RAD 320 biological irradiator. Donor cells (5 × 10^5^) and radiation protector cells (5 × 10^5^) isolated from BM were mixed in 1× PBS and transplanted into the recipient mice through retro-orbital injection. The mice were fed with water supplemented with trimethoprim/sulfamethoxazole for 4 weeks after transplantation.

### FACS analysis

For FACS analysis, cells were stained with antibodies in staining buffer (1× PBS, 2% FBS) and incubated at 4°C for 30 minutes. The samples were washed once with staining buffer before subjected to FACS analysis with the use of a BD LSRII. The antibodies used in this study include anti–Mac-1(eBioscience), anti–Gr-1(eBioscience), anti–c-kit (eBioscience).

### Western Blot analysis

Total proteins were extracted by RIPA buffer and separated by electrophoresis by 8-10% PAGE gel. The protein was transferred to the nitrocellulose membrane and blotted with primary antibodies. The antibodies used for Western Blot and IP-Western Blot analyses included: rabbit anti-Flag (1:1000, cell signaling) and IRDye 680 donkey anti-rabbit second antibody (1: 10000, Li-Cor). The images were developed by Odyssey Li-Cor Imager (Li-Cor).

### Quantitative RT-PCR and ChIP-qPCR assays

RNA was extracted and purified from cells with the use of Qiashredder (QIAGEN) and RNeasy (QIAGEN) spin columns. Total RNA (1 μg) was subjected to reverse transcription using Iscript reverse transcription supermix (Bio-Rad). cDNA levels were assayed by real-time PCR using iTaq universal SYBR green supermix (Bio-Rad) and detected by CFX386 Touch Real-Time PCR detection system (Bio-Rad). Primer sequences for qPCR are listed in Supplementary Table 5 The expression of individual genes is normalized to expression level of *Gapdh*. ChIP assays that used rabbit anti-ASH1L antibody (in house), rabbit anti-H3K36me2 antibody (Abcam), rabbit anti-MLL-AF9-Flag (Cell signaling) were carried out according to the previously reported protocol with the following modifications: ~2 ug antibodies were used in the immunoprecipitation, and chromatin-bound beads were washed 3 times each with TSEI, TSEII, and TESIII followed by 2 washes in 10mM Tris, pH 7.5, 1mM EDTA. Histone modification ChIPs were carried out as previously reported. DNA that underwent ChIP was analyzed by quantitative PCR (qPCR), and data are presented as the percentage of input as determined with CFX manager 3.1 software. Primers for qPCR and ChIP assays are listed in supplementary Tables S3, respectively.

### RNA-seq sample preparation for HiSeq4000 sequencing

RNA was extracted and purified from cells using QI shredder (Qiagen) and RNeasy (Qiagen) spin columns. Total RNA (1 μg) was used to generate RNA-seq library using NEBNext Ultra Directional RNA library Prep Kit for Illumina (New England BioLabs, Inc) according to the manufacturer’s instructions. Adapter-ligated cDNA was amplified by PCR and followed by size selection using agarose gel electrophoresis. The DNA was purified using Qiaquick gel extraction kit (Qiagen) and quantified both with an Agilent Bioanalyzer and Invitrogen Qubit. The libraries were diluted to a working concentration of 10nM prior to sequencing. Sequencing on an Illumina HiSeq4000 instrument was carried out by the Genomics Core Facility at Michigan State University.

### RNA-seq data analysis

RNA-Seq data analysis was performed essentially as described previously. All sequencing reads were mapped mm9 of the mouse genome using Tophat2[25]. The mapped reads were normalized to reads as Reads Per Kilobase of transcript per Million mapped reads (RPKM). The differential gene expression was calculated by Cuffdiff program and the statistic cutoff for identification of differential gene expression is p < 0.01 and 1.5-fold RPKM change between samples[26]. The heatmap and plot of gene expression were generated using plotHeatmap and plotProfile in the deepTools program[27]. The differential expressed gene lists were input into the David Functional Annotation Bioinformatics Microarray Analysis for the GO enrichment analyses (https://david.ncifcrf.gov/).

### Statistical analysis

All statistical analyses were performed using GraphPad Prism 8 (GraphPad Software). Parametric data were analyzed by a two-tailed *t* test or two-way ANOVA test for comparisons of multiple samples. *P* values < 0.05 were considered statistically significant. Data are presented as mean ± SEM.

## Results

### ASH1L is required for the initiation of MLL-AF9-induced leukemic transformation *in vitro*

To examine the function of ASH1L in MLLr-induced leukemogenesis, we generated an *Ash1L* conditional knockout (*Ash1L*-cKO) mouse line in which two *LoxP* elements inserted into the exon 4 flanking regions[24]. A CRE recombinase-mediated deletion of exon 4 resulted in altered splicing of mRNA that created a premature stop codon before the sequences encoding the first functional AWS (*<u>A</u>*ssociated *<u>W</u>*ith *<u>S</u>*ET) domain (Fig. S1a, S1b). The *Ash1L*-cKO mice were further crossed with the Rosa26-CreER^T2^ mice to generate a tamoxifen-inducible *Ash1L* knockout line (*Ash1L*^2f/2f^;Rosa26-CreER^T2^), which allowed us to study the function of ASH1L in leukemogenesis *in vitro* and *in vivo*.

Using this *Ash1L*-cKO mouse model, we set out to investigate the role of ASH1L in the initiation of MLL-AF9-induced leukemic transformation. To this end, we isolated the bone marrow cells from wild-type (*Ash1L*^+/+^;*Cre-ER^T2^*) and *Ash1L*-cKO (*Ash1L*^2f/2f^;*Cre-ER^T2^*) mice, respectively. The c-kit^+^ HPCs were further enriched by the c-kit antibody-conjugated magnetic beads (Fig. S1c). The HPCs were cultured in the HPC medium supplemented with murine IL-3, IL-6, and SCF for three days, and followed by transduction of retroviral vectors expressing a *MLL1-AF9* fusion gene or control empty viruses (EV). After transduction, the cells were cultured in the suspension medium with 4-hydroxytamoxifen (4-OHT) for five days to induce *Ash1L* gene deletion in the *Ash1L*-cKO HPCs (Fig. 1a). The quantitative RT-PCR (qRT-PCR) analysis showed that the *Ash1L* expression reduced to less than 5% at the mRNA level in the *Ash1L*-deleted cells (Fig. 1b). To investigate the effect of *Ash1L* loss on the initiation of MLL-AF9-induced leukemic transformation *in vitro*, we performed the serial colony replating assays by plating the cells on the semi-solid methylcellulose medium to examine the leukemic transformation. The results showed that although the cells transduced with *MLL-AF9* or empty vectors had comparable colony numbers in the first round of plating, the cells transduced with control empty vectors did not form colonies in the following rounds of replating. In contrast, both wild-type and *Ash1L*-cKO HPCs transduced with *MLL-AF9* retroviruses formed colonies in all three rounds of plating, indicating the successful leukemic transformation by the *MLL-AF9* transgene *in vitro*. Notably, compared to the MLL-AF9-transduced wild-type cells, the *Ash1L*-deleted cells had reduced colony numbers in the second and third rounds of plating, suggesting that loss of *Ash1L* in HPCs compromised the MLL-AF9-induced leukemic transformation (Fig. 1c, d). Further qRT-PCR analyses showed that the MLL-transformed wild-type and *Ash1L*-cKO cells had the comparable expression levels of *MLL-AF9* transgene, suggesting that the difference in colony numbers was not due to a disparity in *MLL-AF9* transgene gene expression (Fig. S1d).

**Figure 1.**
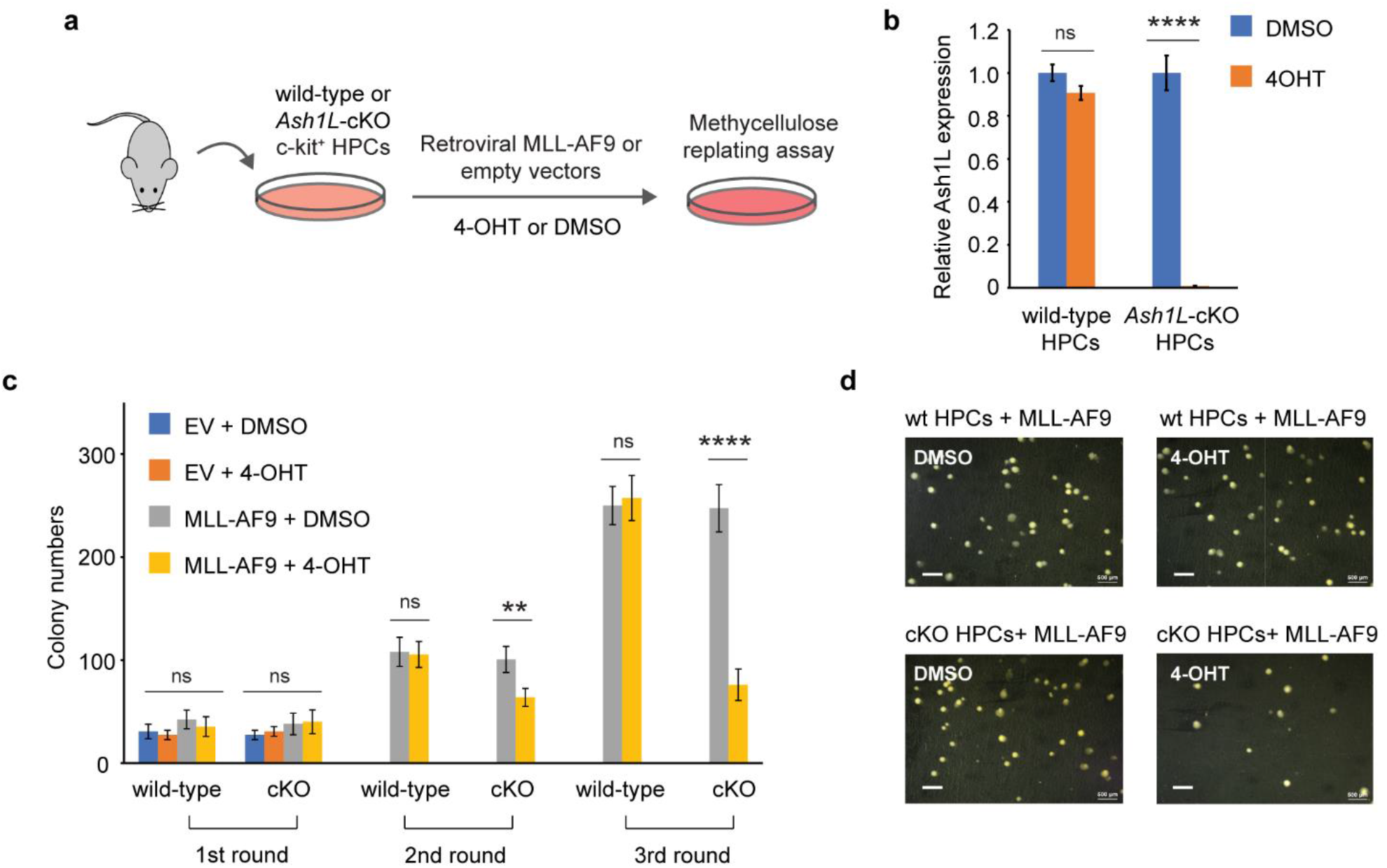
ASH1L is required for the initiation of MLL-AF9-induced leukemic transformation. (a) Schematic experimental procedure. (b) qRT-PCR analysis showing the *Ash1L* expression levels in wild-type and *Ash1L*-cKO cells after treated with 4-OHT or DMSO. The results were normalized against levels of *Gapdh* and the expression level in DMSO-treated cells was arbitrarily set to 1. The error bars represent mean ± SEM, n = 3 per group. \**P* < 0.05; \*\**P* < 0.01; \*\*\**P* < 0.001, ns, not significant. (c) Methylcellulose replating assays showing the colony numbers for each round of plating. The error bars represent mean ± SEM, n = 3 per group. \*\**P* < 0.01; \*\*\*\**P* < 0.0001, ns, not significant. (d) Photos showing the representative colony formation on methylcellulose plates for each group. Bar = 0.5 mm.

### ASH1L is required for the maintenance of MLL-AF9-induced leukemic cells *in vitro*

Next, we set out to examine whether *Ash1L* was required for the maintenance of MLL-AF9 transformed cells. To this end, we transduced both wild-type and *Ash1L*-cKO HPCs with *MLL-AF9* retroviruses and plated the transduced cells onto the methylcellulose medium. After three rounds of replating, the transformed colonies were manually picked and cultured in the suspension medium supplemented with 4-OHT for 5 days to induce deletion of *Ash1L* in the *Ash1L*-cKO cells. The cells were further maintained in suspension culture without 4-OHT for 5 days before plated onto the methylcellulose to examine the colony forming capacity (Fig. 2a). The results showed that compared to the wild-type MLL-AF9-transformed cells, the *Ash1L*-deleted cells had marked reduced colony formation (Fig. 2b, c), suggesting that ASH1L was required for the maintenance of MLL-AF9 transformed cells *in vitro*.

**Figure 2.**
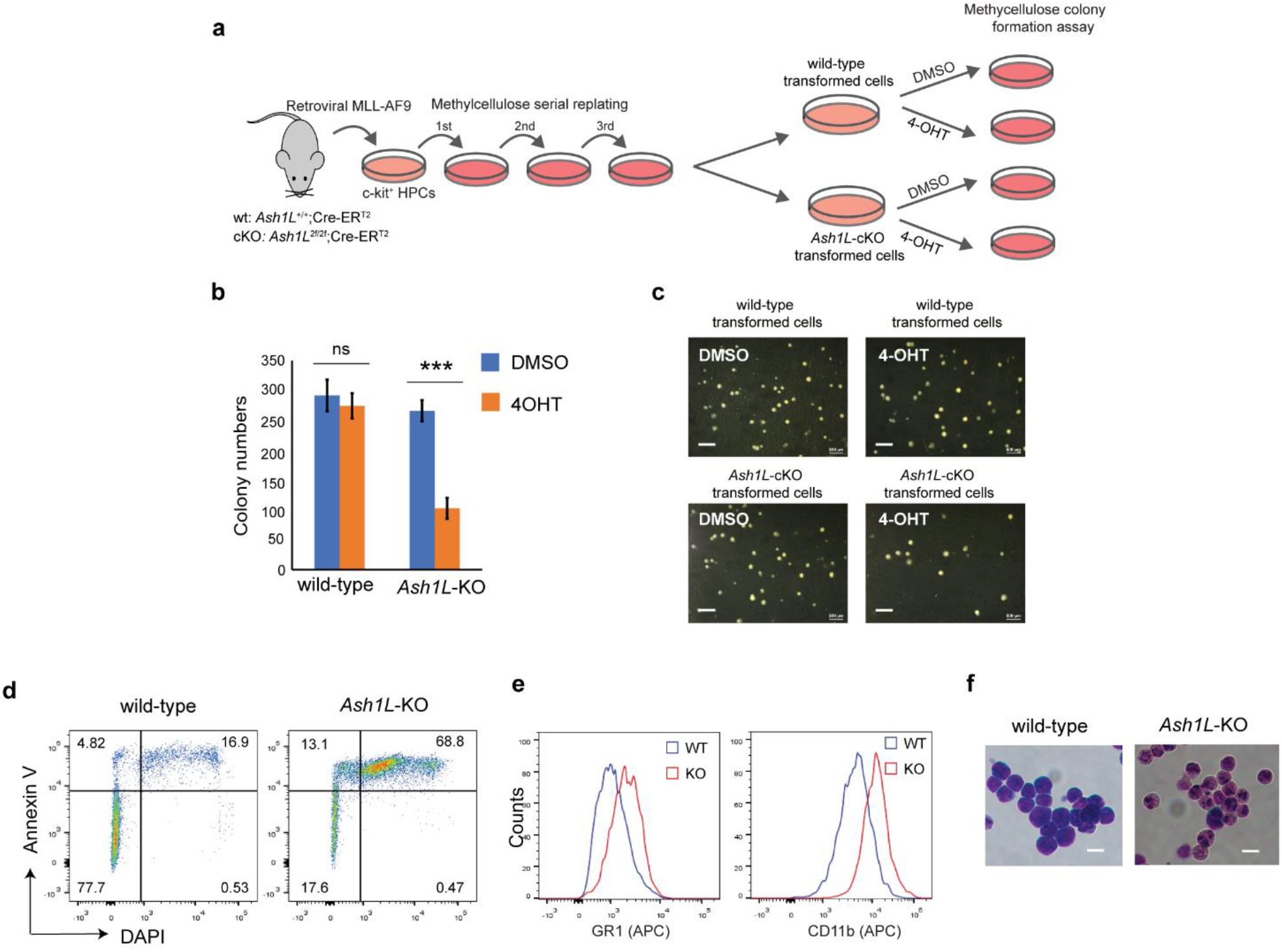
ASH1L is required for the maintenance of MLL-AF9-induced leukemic cells *in vitro*. (a) Schematic experimental procedure. (b) Methylcellulose colony formation assays showing the colony numbers. The error bars represent mean ± SEM, n = 3 per group. \*\*\**P* < 0.001; ns, not significant. (c) Photos showing the representative colony formation on methylcellulose plates for each group. Bar = 0.5 mm. (d) Representative FACS results showing the Annexin V+ and DAPI+ populations of wild-type and *Ash1L*-KO MLL-AF9-transformed cells. (e) Representative FACS results showing the GR1 and CD11b expression of wild-type and *Ash1L*-KO MLL-AF9-transformed cells. (f) Photos showing the Wright-Giemsa staining of wild-type and *Ash1L*-KO MLL-AF9-transformed cells. Bar = 10 μm.

To examine the cellular responses to the *Ash1L* depletion, we performed the FACS analysis to examine the cell death in response to the loss of *Ash1L* in the MLL-AF9-transformed cells. The results showed that compared to the wild-type cells, the *Ash1L*-deleted cells had increased populations of both early apoptotic cells (Annexin V+/DAPI-) and late dead cells (Annexin V+/DAPI+) (Fig. 2d), suggesting that the loss of *Ash1L* induced cell death of MLL-AF9-transformed cells. Moreover, FACS analyses showed that compared to the wild-type transformed cells, the *Ash1L*-deleted cells had increased expression of myeloid differentiation surface markers CD11b and GR-1 (Fig. 2e). Morphologically, the wild-type transformed cells displayed leukoblast-like morphology with enlarged dark stained nuclei, while the *Ash1L*-deleted cells had light-stained and segmented nuclei, a feature indicating the differentiation towards matured myeloid cells (Fig. 2f). Taken together, these results suggested that ASH1L was required for the maintenance of MLL-AF9-transformed cells through suppressing cell death and differentiation.

### ASH1L is required for the MLL-AF9-induced leukemia development *in vivo*

To determine whether ASH1L was required for the MLL-AF9-induced leukemogenesis *in vivo*, we performed leukemia transplantation assays and monitor the leukemia development in the recipient mice. To this end, the wild-type and *Ash1L*-deleted MLL-AF9-transformed cells were labeled with GFP by transduction with lentiviral-GFP vectors, mixed with normal protective bone marrow cells, and transplanted into the total-body-irradiated (TBI) syngeneic recipient mice (Fig. 3a). Four weeks after transplantation, FACS analysis showed that the mice transplanted with wild-type leukemic cells had higher GFP+ leukemic cell populations in the peripheral blood compared to the mice received with *Ash1L*-KO leukemic cells (Fig. 3b), which was consistent with the higher leukemic cell numbers in the peripheral blood smears and splenomegaly found in the mice transplanted with wild-type leukemic cells (Fig. 3c, d). All mice transplanted with wild-type leukemic cells died within 3 months after transplantation, and the median survival time was around 9 weeks. In contrast, the median survival time for the mice transplanted with *Ash1L*-deleted cells was around 12.5 weeks and the overall survival time was 4 weeks longer than the mice transplanted with wild-type leukemic cells (Fig. 3e). These results suggested that ASH1L in the MLL-AF9-transformed leukemic cells was required for the development and progression of leukemia *in vivo*.

**Figure 3.**
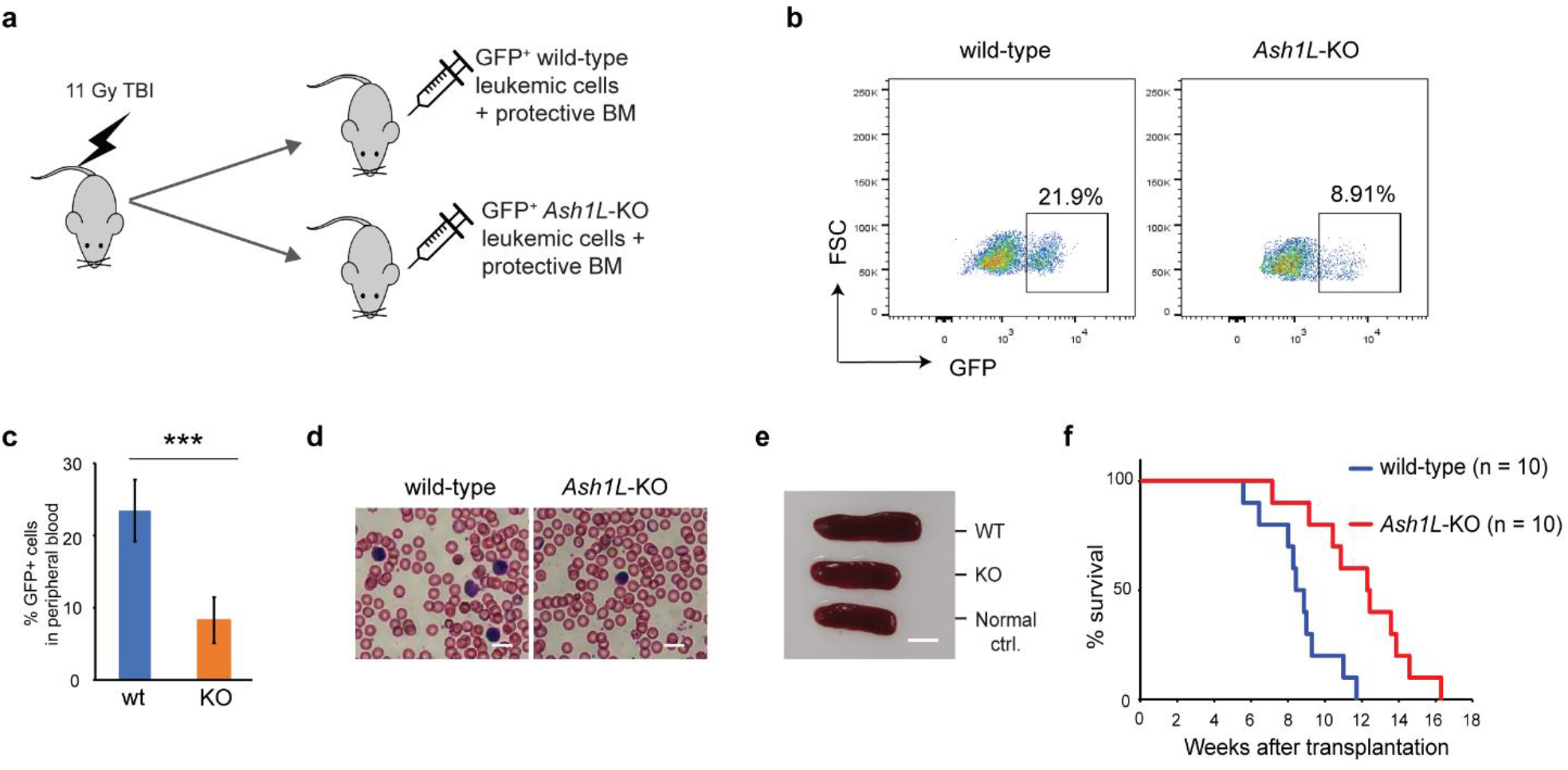
ASH1L is required for the MLL-AF9-induced leukemia development *in vivo*. (a) Schematic experimental procedure. (b) Representative FACS analysis showing the GFP+ leukemic cell populations in the peripheral blood of mice transplanted with wild-type or *Ash1L*-KO MLL-AF9-transformed cells. The error bars represent mean ± SEM, n = 3 per group. \*\*\**P* < 0.001; ns, not significant. (c) Quantitative results showing the percentage of GFP+ leukemic cell populations in the peripheral blood of mice transplanted with wild-type or *Ash1L*-KO MLL-AF9-transformed cells. The error bars represent mean ± SEM, n = 5 per group. \*\**P* < 0.01. (d) Photos showing the leukemic cells in the peripheral blood smear of mice transplanted with wild-type or *Ash1L*-KO MLL-AF9-transformed cells. (e) Photos showing the representative spleen size from the normal control mice (Normal ctrl.), mice transplanted with wild-type (WT) or *Ash1L*-KO (KO) MLL-AF9-transformed cells. (f) The survival curve of mice transplanted with wild-type or *Ash1L*-KO MLL-AF9-transformed cells. n = 15 mice per group.

### The enzymatic activity of ASH1L is required for its function in promoting MLL-AF9-induced leukemic transformation

Next, we set out to determine whether the histone methyltransferase activity of ASH1L was required for its function in promoting MLL-AF9-induced leukemic transformation. To this end, the *Ash1L*-cKO HPCs were infected with retroviruses expressing *MLL-AF9* transgene, followed by transduced with lentiviral vectors expressing either wild-type ASH1L or catalytic-dead mutant ASH1L(H2214A)[21]. The transformed cells were treated with 4-OHT to induce deletion of endogenous *Ash1L* gene (Fig. 4a). WB analysis showed that both wild-type and mutant exogenous ASH1L had a similar expression level (Fig. 4b). The cells were further plated onto the methylcellulose medium to examine the colony formation (Fig. 4a). The results showed that compared to the wild-type ASH1L expressed cells, the cells with ectopic expression of catalytic-dead mutant ASH1L had reduced colony formation (Fig 4c). Similar to the *Ash1L*-deleted cells, the *Ash1L*-deleted cells rescued with mutant ASH1L had increased cell death and upregulated expression of myeloid differentiation markers of CD11b and GR-1 (Fig. 4e, f). These results suggested that ASH1L histone methyltransferase activity was required for its function in promoting MLL-AF9-induced leukemogenesis by inhibiting cell death and blocking myeloid differentiation.

**Figure 4.**
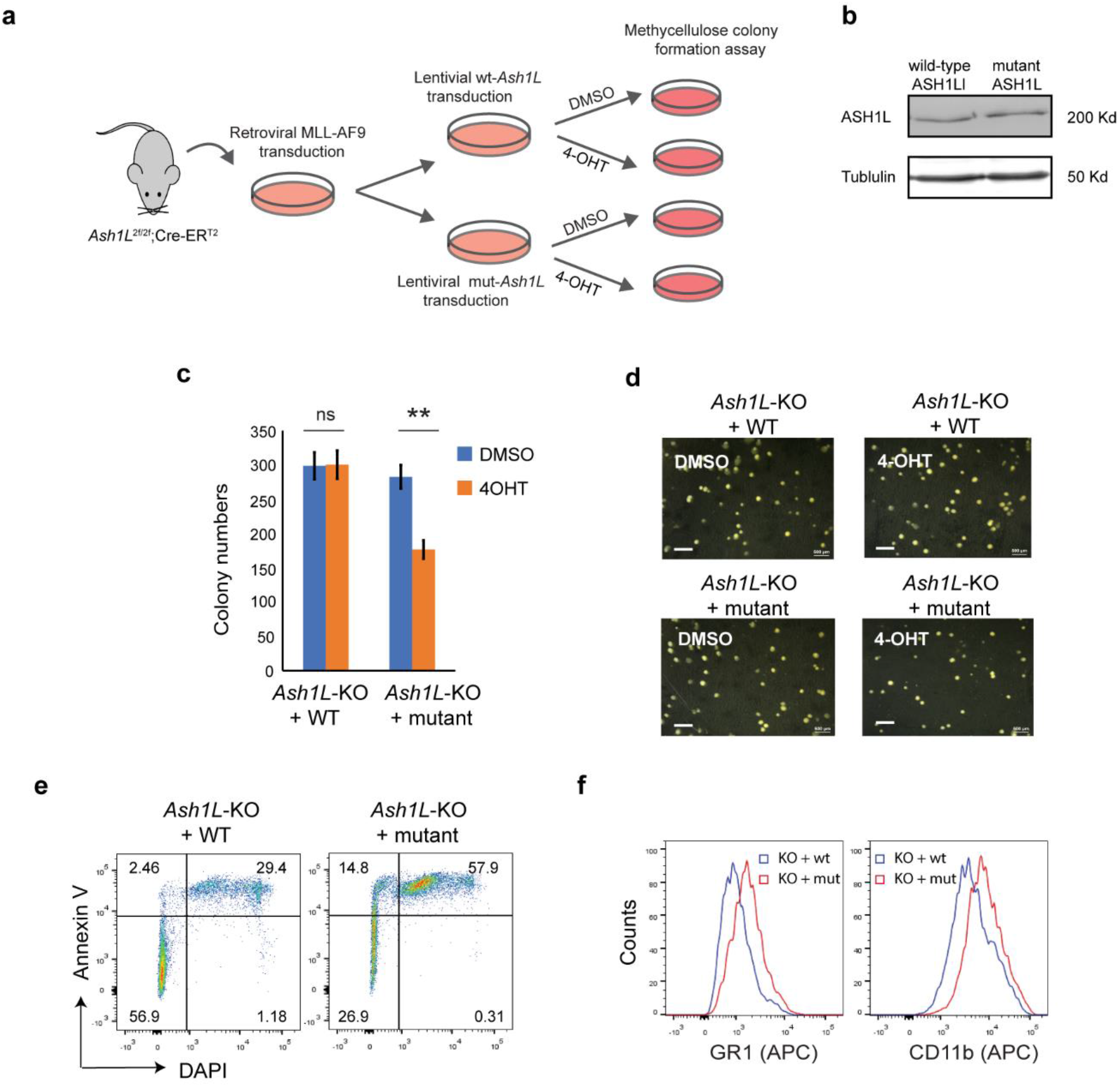
The enzymatic activity of ASH1L is required for its function in promoting MLL-AF9-induced leukemic transformation. (a) Schematic experimental procedure. (b) WB analysis showing the ectopic expression of wild-type and mutant ASH1L. (c) Methylcellulose colony formation assays showing the colony numbers. The error bars represent mean ± SEM, n = 3 per group. \*\**P* < 0.01; ns, not significant. (d) Photos showing the representative colony formation on methylcellulose plates for each group. Bar = 0.5 mm. (e) Representative FACS results showing the Annexin V+ and DAPI+ populations of *Ash1L*-KO cells rescued with wild-type and mutant ASH1L. (f) Representative FACS results showing the GR1 and CD11b expression of *Ash1L*-KO cells rescued with wild-type and mutant ASH1L.

### ASH1L facilitates the MLL-AF9-induced leukemogenic gene expression

To examine the molecular mechanisms underlying the function of ASH1L in promoting MLL-AF9-induced leukemogenesis, we performed the RNA-seq analysis to examine the transcriptome changes in normal HPCs, wild-type and *Ash1L*-deleted MLL-AF9-tranformed cells. The results showed that compared to normal HPCs, the MLL-AF9-transformed cells had 1,021 upregulated and 1,228 downregulated genes (cutoff: fold changes > 1.5, FDR < 0.05), respectively (Fig. 5a). The gene ontology (GO) enrichment analysis showed that both upregulated and downregulated genes were involved in immune processes and inflammatory responses (cutoff: FDR < 0.05) (Tables S1 and S2), suggesting that MLL-AF9 fusion proteins disrupted the normal differentiation and mis-regulated the normal function of myeloid cells. Notably, multiple genes, such as *Hoxa5*, *Hoxa7*, *Hoxa9*, *Hoxa10* and *MeisI* that were known to mediate the MLL-AF9-induced leukemogenesis, were highly expressed in the MLL-AF9-transformed cells (Fig. 5b). Further RNA-seq analysis showed that compared to MLL-AF9-transformed wild-type cells, the *Ash1L*-deleted cells had 372 upregulated gene and 472 downregulated genes (cutoff: fold changes > 1.5, FDR < 0.05), respectively (Fig. 5c). Cross-examining these two data sets revealed that 105 genes, including *Hoxa5*, *Hoxa7*, *Hoxa9*, *Hoxa10*, and *MeisI* that were highly expressed in the wild-type MLL-AF9-transformed cells, were downregulated in the *Ash1L*-deleted cells (Fig. 5d, e). Overall, these results suggested that ASH1L promoted the MLL-AF9-induced leukemogenesis by facilitating the MLL-AF9-induced leukemic gene expression.

**Figure 5.**
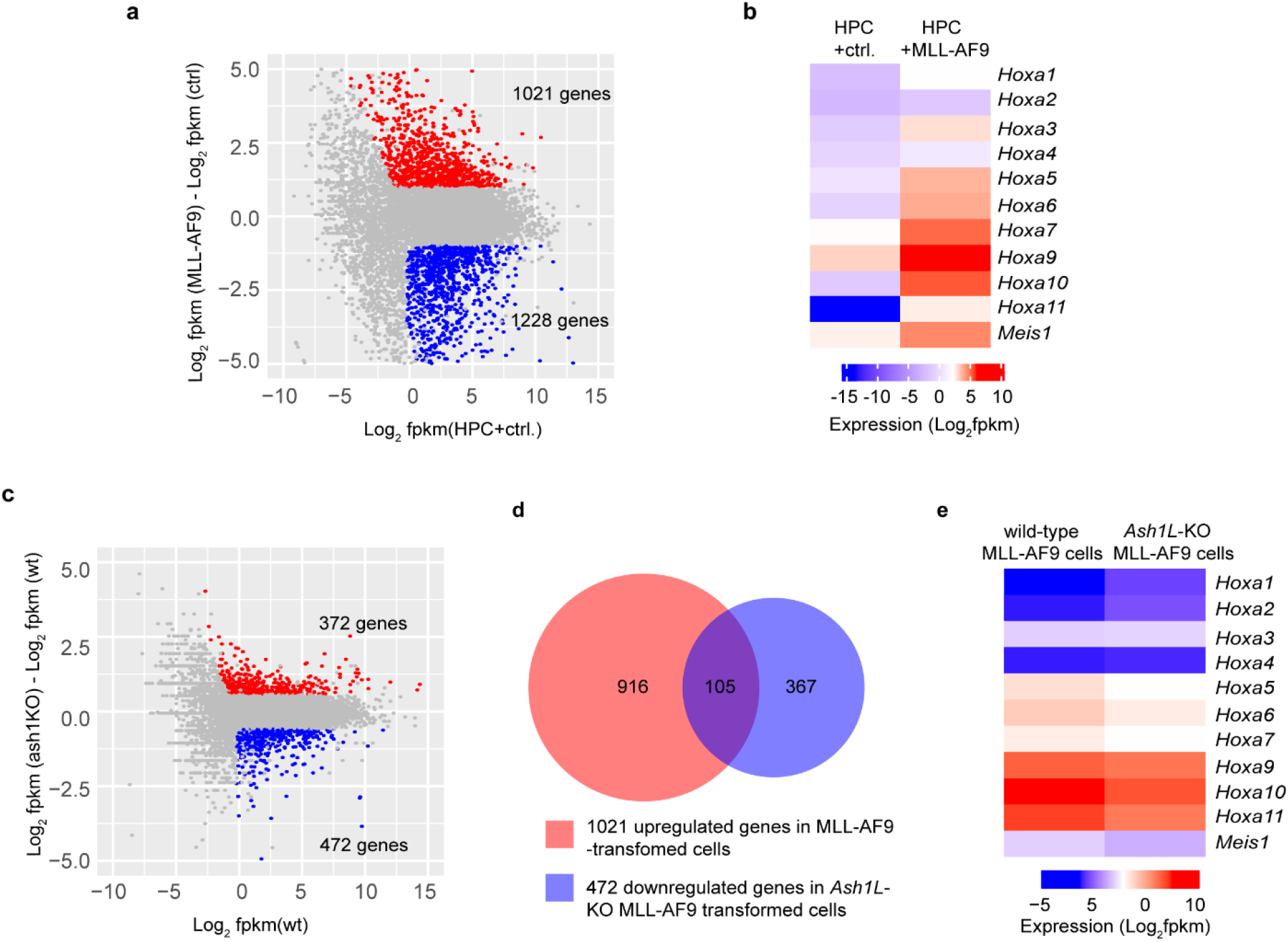
ASH1L facilitates the MLL-AF9-induced leukemogenic gene expression. (a) Plot showing 1021 up- and 1228 down-regulated genes in the MLL-AF9-transformed cells compared to the normal HPCs. (b) Heatmap showing the upregulation of *Hoxa* gene cluster and *MeisI* in the MLL-AF9-transformed cells compared to normal HPCs. (c) Plot showing 372 up- and 472 down-regulated genes in the *Ash1L*-KO MLL-AF9-transformed cells compared to the wild-type MLL-AF9-transformed cells. (d) Venn diagram showing the 105 genes upregulated in the MLL-AF9-transformed cells and downregulated in the *Ash1L*-KO cells. (e) Heatmap showing the *Hoxa* gene cluster and *MeisI* downregulated in the *Ash1L*-KO cells compared to the wild-type MLL-AF9-transformed cells.

### ASH1L binds and mediates the histone H3K36me2 modification at *Hoxa9* and *Hoxa10* gene promoters

To determine whether ASH1L directly regulated the expression of MLL-AF9 target genes, we performed chromatin immunoprecipitation (ChIP) coupled with quantitative PCR (ChIP-qPCR) assays to examine the ASH1L occupancy, MLL-AF9 occupancy, and histone H3K36me2 modification at the gene promoters, transcriptional starting sites (TSS), transcriptional ending sites (TES) of *Hoxa9* and *Hoxa10*, two MLL-AF9 target genes that were shown to be activated in the wild-type transformed cells and have reduced expression in the *Ash1L*-deleted cells (Fig. 5b, e). The results showed that both ASH1L occupancy and histone H3K36me2 were enriched at the *Hoxa9* and *Hoxa10* promoters compared to that on the TES and the long terminal repeat (LTR) of endogenous retroviruses. Furthermore, compared to wild-type MLL-AF9-transformed cells, both ASH1L occupancy and histone H3K36me2 modification were reduced at the gene promoters in the *Ash1L*-deleted cells, suggesting that ASH1L bound to the *Hoxa9* and *Hoxa10* gene promoters directly and mediated local histone H3K36me2 modification. However, the MLL-AF9 occupancy at both gene promoters did not show significant difference between wild-type and *Ash1L*-deleted MLL-AF9-transformed cells, suggesting the ASH1L-mediated histone H3K36me2 did not to affect the binding of MLL-AF9 fusion protein to the gene promoters.

## Discussion

Chromosomal 11q23 translocations generate various MLL fusion proteins that contain the *N*-terminal portion of MLL1 and different fusion partners including AF9[28, 29]. Previous studies have demonstrated that the *N*-terminal MLL1 is critical for the recruitment of MLL fusion proteins to chromatin through its CxxC-zinc finger (CxxC-zf) domain as well as its interacting proteins MENIN and LEDGF, while the *C*-terminal fusion partners interact with multiple trans-activators to induce transcriptional activation[16]. Since the MLL fusion proteins lose the MLL1 *C*-terminal SET domain and its-associated histone H3K4 methyltransferase activity, it is unclear whether other histone KMTase-mediated histone modifications are required for the MLL fusion proteins to activate leukemogenic gene expression and induce leukemia development.

ASH1L is another member of TrxG proteins that facilitate transcriptional activation[8]. Biochemically, ASH1L is a histone KMT mediating histone H3K36me2 modification[20]. Recent studies reported that ASH1L and MLL1 co-occupied the same gene promoters to activate gene expression, suggesting ASH1L and MLL function synergistically in activating gene expression in normal development and leukemogenesis[19, 21–23]. However, the functional roles of ASH1L and its-mediated histone H3K36me2 in the MLLr leukemia development have not been addressed using *Ash1L* gene knockout animal models.

In this study, we used an *Ash1L* conditional knockout mouse model to show that ASH1L and its histone methyltransferase activity are required for the MLL-AF9-induced leukemogenesis. First, genetic deletion of ASH1L in normal HPCs largely impairs the MLL-AF9-induced colony formation in serial methylcellulose replating assays (Fig. 1), suggesting ASH1L is required for the initiation of MLL-AF9-induced leukemic transformation. Second, loss of ASH1L in the MLL-AF9-transformed cells largely impaired the colony formation *in vitro* and delayed the leukemia development in the recipient mice transplanted with leukemic cells (Figs. 2 and 3), suggesting ASH1L is required for the maintenance of MLL-AF9-transformed cells *in vitro* and leukemia progression *in vivo*. Importantly, the impaired ASH1L’s function in the *Ash1L*-KO cells could be rescued by the wild-type but not the catalytic-dead mutant ASH1L (Fig. 4), suggesting that the histone methyltransferase activity is required for its function in promoting MLL-AF9-induced leukemogenic transformation, which is consistent with a recent study showing that the SET domain is required for the MLL-AF9-induced leukemic transformation[30].

At the cellular level, we observed that the loss of ASH1L in MLL-AF9-transformed cells induced cell death and myeloid differentiation, which could be rescued by the wild-type but not the catalytic-dead mutant ASH1L (Figs. 2 and 4), suggesting that ASH1L promotes MLL-AF9-induced leukemic transformation though inhibit cell apoptosis and block cell differentiation. These results are consistent with the molecular findings that ASH1L is required for the full activation of MLL-AF9 target genes including *Hoxa* gene cluster and *MeisI* (Fig. 5), which were known to play important roles in leukemogenesis through inhibiting cell death and blocking normal cell differentiation[31–33]. Finally, the ChIP assays showed that both ASH1L occupancy and histone H3K36me2 modification were enriched at the promoters of MLL-AF9 target genes *Hoxa9* and *Hoxa10* in the wild-type transformed cells (Fig. 6), indicating the ASH1L regulates the MLL-AF9 target genes through directly chromatin binding and its-mediated histone H3K36me2 modification.

**Figure 6.**
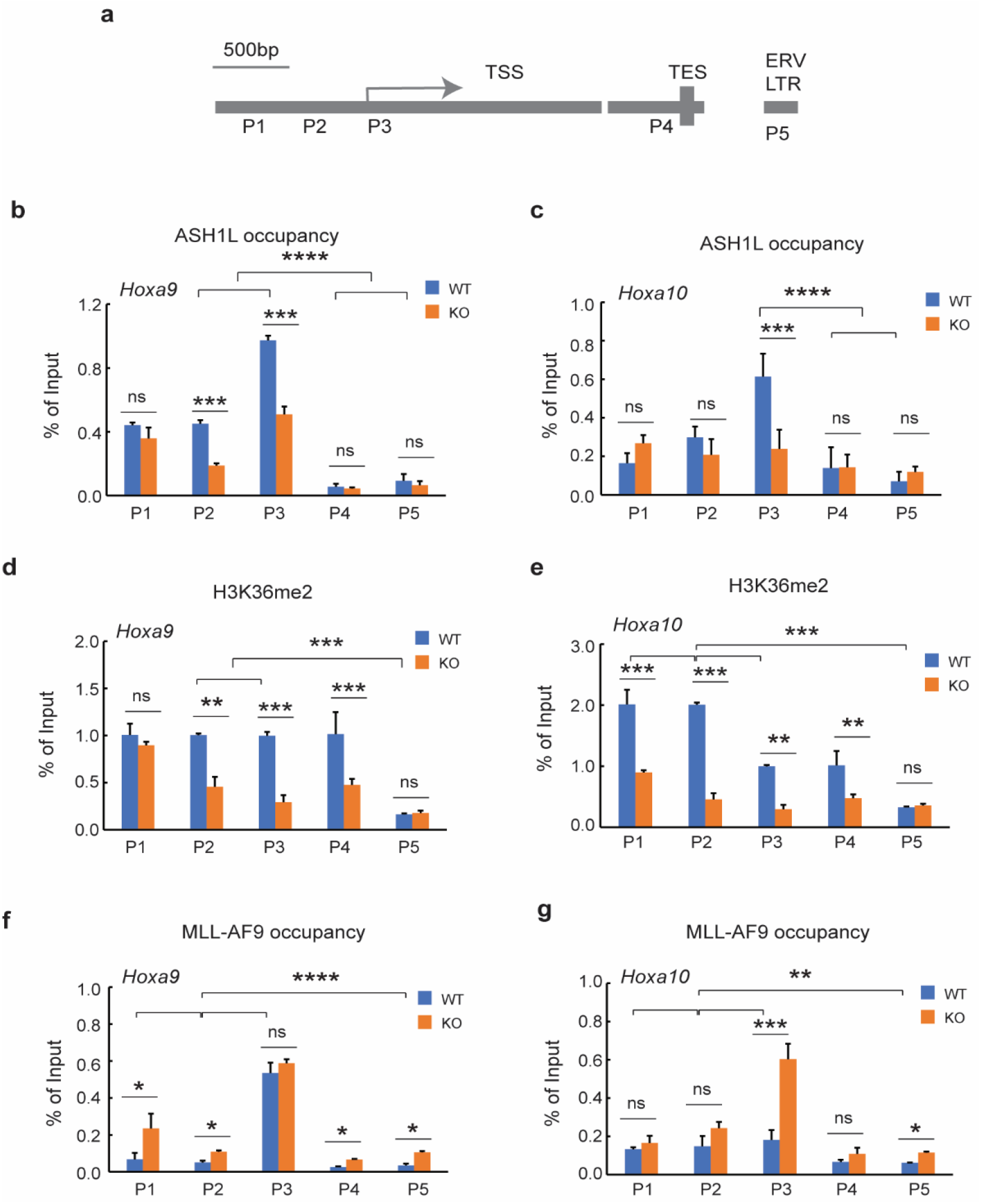
ASH1L binds and mediates histone H3K36me2 modification at *Hoxa9* and *Hoxa10* gene promoters. (a) Plot showing the locations of ChIP-qPCR amplicons at the *Hoxa9* and *Hoxa10* gene loci and LTR of endogenous retrovirus. (b-c) ChIP-qPCR analysis showing the ASH1L occupancy at *Hoxa9* and *Hoxa10* gene loci in the wild-type and *Ash1L*-KO MLL-AF9-transformed cells. (d-e) ChIP-qPCR analysis showing the histone H3K36me2 at *Hoxa9* and *Hoxa10* gene loci in the wild-type and *Ash1L*-KO MLL-AF9-transformed cells. (f-g) ChIP-qPCR analysis showing the MLL-AF9 occupancy at *Hoxa9* and *Hoxa10* gene loci in the wild-type and *Ash1L*-KO MLL-AF9-transformed cells. Note: for Figs. b-e, the error bars represent mean ± SEM, n = 3 biological replicates. \**P* < 0.05, \*\**P* < 0.01, \*\*\**P* < 0.001; \*\*\*\**P* < 0.0001; ns, not significant.

Previous studies have shown that the PWWP domain of LEDGF is required for the recruitment of MLL fusion proteins through its binding to histone H3K36me2[13, 15, 16]. However, our ChIP analysis did not reveal the reduction of MLL-AF9 occupancy at the *Hoxa9* and *Hoxa10* promoters in the *Ash1L*-KO cells (Fig. 6f, g), suggesting the MLL-AF9 fusion protein could bind to its target regions though other recruiting mechanisms, such as through the CxxC-zf domain binding to the unmethylated CpG-rich promoters[34], and the reduced H3K36me2 at gene promoters in the *Ash1L*-KO cells might impair the *Hoxa* gene expression through mechanisms other than the recruitment of MLL-AF9 fusion protein.

In summary, our study reveals that the histone H3K36me2-specific methyltransferase ASH1L and its enzymatic activity play a critical role in promoting MLL-AF9-induced leukemogenesis, which provides an important molecular basis for targeting ASH1L and its enzymatic activity to treat MLL-r leukemias.

## Data availability statement

The RNA-seq data presented in this study is being deposited to the Gene Expression Omnibus database. Other data have been disclosed in the article/Supplementary Material.

## Author contributions

J.H. conceived the project. M.B.A., Y.G., Y.W. and J.H. performed the experiments. J.H. and G.I.M. performed the sequencing data analysis. J.H. interpreted the data and wrote the manuscript.

## Acknowledgements

MSU genomics core facility processed the next-generation sequencing.

